# FleN contributes to heterogeneous swimming at high temperatures in *Pseudomonas syringae*

**DOI:** 10.1101/372060

**Authors:** Kevin L. Hockett, Steven E. Lindow

**Affiliations:** Department of Plant and Microbial Biology, University of California, Berkeley, California, USA

**Keywords:** *Pseudomonas*, Motility, Thermoregulation, Flagellum, Bacterial Heterogeneity

## Abstract

Motility is generally conserved among many animal and plant pathogens. Environmental conditions, however, significantly impact expression of the motile phenotype. In this study, we describe a novel heterogeneous motility phenotype in *Pseudomonas syringae*, where under normally suppressive incubation conditions (30°C) punctate colonies arise that are spatially isolated from the point of inoculation, giving rise to a motility pattern we term constellation swimming (CS). We demonstrate that this phenotype is reproducible, reversible, and dependent on a functioning flagellum. Mirroring the heterogeneous motility phenotype, we demonstrate the existence of a sub-population of cells under non-permissive conditions that express flagellin (*fliC*) at levels similar to cells incubated under permissive conditions using both quantitative single cell microscopy and flow cytometry. To understand the genetics underlying the CS phenotype, we selected for naturally arising mutants that exhibited a normal swimming phenotype at the warmer incubation temperature. Sequencing these mutants recovered several independent non-synonymous mutations within FleN (also known as FlhG) as well as mutations within the promoter region of FleQ, the master flagellum regulator in *Pseudomonas*. We further show that nutrient depletion is the likely underlying cause of CS, as reduced nutrients will stimulate both *fliC* expression and a normal swimming phenotype at 30 °C.

## INTRODUCTION

Flagellar-mediated motility is a trait utilized by bacteria inhabiting diverse environments, which allows cells to seek out favorable (nutrient replete) locations. Such motility is regulated by diverse environmental factors, including temperature, carbon source, growth phase, pH, and osmolarity (reviewed in (Soutourina & Bertin, 2003)), all of which affect flagellar synthesis. This tight control likely reflects the substantial cellular resources required for flagellum synthesis (Macnab, 1996); bacteria would be expected to limit production only to circumstances where motility would be beneficial. Additionally, synthesis of the flagellum is itself a risky endeavor, leading to cell death in a fraction of expressing cells, regardless of the environment (Fontaine, Stewart, & Lindner, 2008).

*Pseudomonas syringae*, as a colonist of leaf surfaces, is exposed to highly variable environmental conditions, which fluctuate both regularly (in a diurnal pattern) as well as irregularly in response to prevailing weather conditions (Hirano & Upper, 2000b; Lindow & Brandl, 2003). In addition, the resources that *P. syringae* exploits on a leaf are highly spatially variable (Leveau & LINDOW, 2001), and the motility of *P. syringae* has thus been found to be an important fitness factor, presumably by enabling the cells to explore the leaf surface in pursuit of such resources and to avoid stressful sites (Haefele & LINDOW, 1987; Hockett, Burch, & Lindow, 2013). As such, it would be expected that *P. syringae* is adapted to integrate multiple, sometimes conflicting, environmental signals in a manner to express motility in a way that would maximize its survival and proliferation in this habitat. One such strategy may include the formation of phenotypically distinct sub-populations, some of which are pre-acclimated to respond to sudden and unpredictable environmental fluctuations, enabling a *bet-hedging* life strategy (Veening, Smits, & Kuipers, 2008). Such strategies have been documented, and theoretical as well as empirical work has documented their benefit to cells (Libby & Rainey, 2011; Lidstrom & Konopka, 2010; Salathé, Van Cleve, & Feldman, 2009).

Regulation of the bacterial flagellum is a complex process that is achieved by a hierarchy of distinct, interacting regulators (Chevance & Hughes, 2008; McCarter, 2006). This regulatory complexity reflects the elaborate nature of the organelle itself, which requires coordinated temporal expression and spatial organization of the component proteins to produce a functional flagellum. For the majority of species, environmental regulation of flagellar synthesis affects expression of the master regulators, *flhDC* or *fleQ*, depending on the organism (McCarter, 2006; Soutourina & Bertin, 2003). In a few cases, such environmental regulation affects check-points downstream of the master regulator (Horne & Pruss, 2006; Wosten et al., 2010). In addition to the core components of flagellar regulation, certain organisms, such as *Pseudomonas* species, employ additional levels of regulation that ensure the proper localization of the flagellum, as well as restrict the number of flagella produced (McCarter, 2004). Proper localization and quantitative regulation in polarly-flagellated bacteria is achieved through the interaction of two genes, *flhF* and *fleN/flhG* that are absent in peritrichously flagellated bacteria having multiple flagella distributed over the cell body (Kusumoto et al., 2008; Murray & Kazmierczak, 2006; Pandza et al., 2000).

In this work we describe a novel heterogeneous swimming phenotype exhibited by a subset of cells of *P. syringae* incubated at warm temperatures (30°C), temperatures that are inhibitory to expression of numerous flagellar genes (Hockett et al., 2013), as well as for normal swimming motility and show that this is due to a transient variation in gene expression within the population of cells. Examination of stable spontaneous mutants that gained the ability to swim at elevated temperatures revealed a role for the polar flagellum-specific regulator FleN. Moreover, we show that locally reduced nutrient concentration is likely cause for the heterogeneous swimming phenotype, as both elevated *fliC* expression and a normal swimming phenotype at 30 °C are exhibited by cells incubated under low nutrient conditions.

## RESULTS

### *P. syringae* displays a constellation swimming phenotype when incubated at 30°C

Previously, we showed that *P. syringae* incubated at 30°C exhibited dramatically less swimming when compared to cooler incubation temperatures (Hockett et al., 2013). Interestingly, in addition to producing a reduced swimming zone, incubation at a warm temperature resulted in a qualitative change in the swimming phenotype. Cells were not uniformly dispersed, but instead formed punctate colonies at various distances from the inoculation stab (Fig. 1). We refer to this phenotype as constellation swimming (CS). Wild type *P. syringae* cells readily formed spatially separated colonies, while a Δ*flgK* mutant, which is impaired in flagellar-mediated motility did not, indicating that the phenotype required a functioning flagellum and thus reflected active motility (Fig. 2A and B). We tested whether this phenotype could be propagated, as well as reversed, if the colonies were transferred to fresh media. When isolated colonies were transferred to fresh swimming media, the culture exhibited a normal swimming phenotype at 20°C but recapitulated the CS phenotype at 30°C (Fig. 2C and D, respectively). These experiments confirm the CS phenotype is a genetically-encoded trait rather than mis-regulated swimming displayed by a few stable mutants.

**Figure 1.**
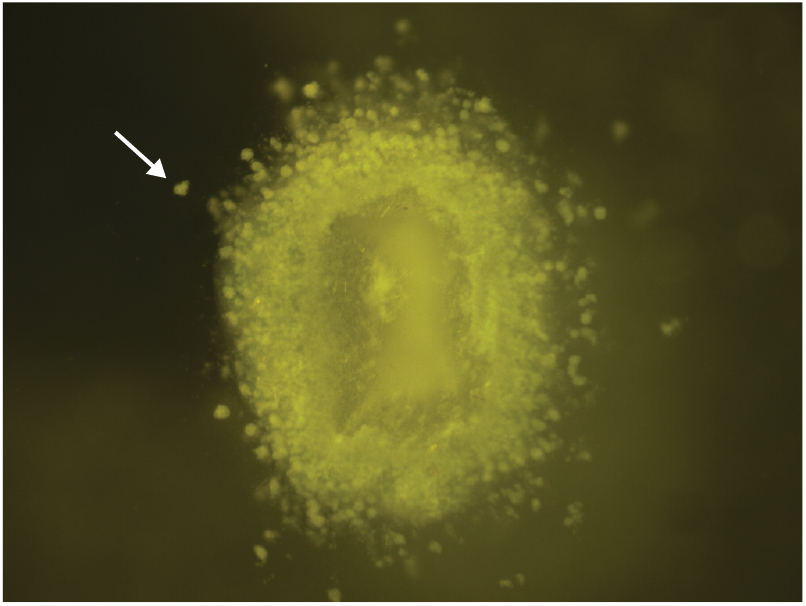
Wild type *Psy* B728a exhibiting constellation swimming (CS) at 30 °C. The white arrow indicates a colony spatially separated from the point of inoculation.

**Figure 2.**
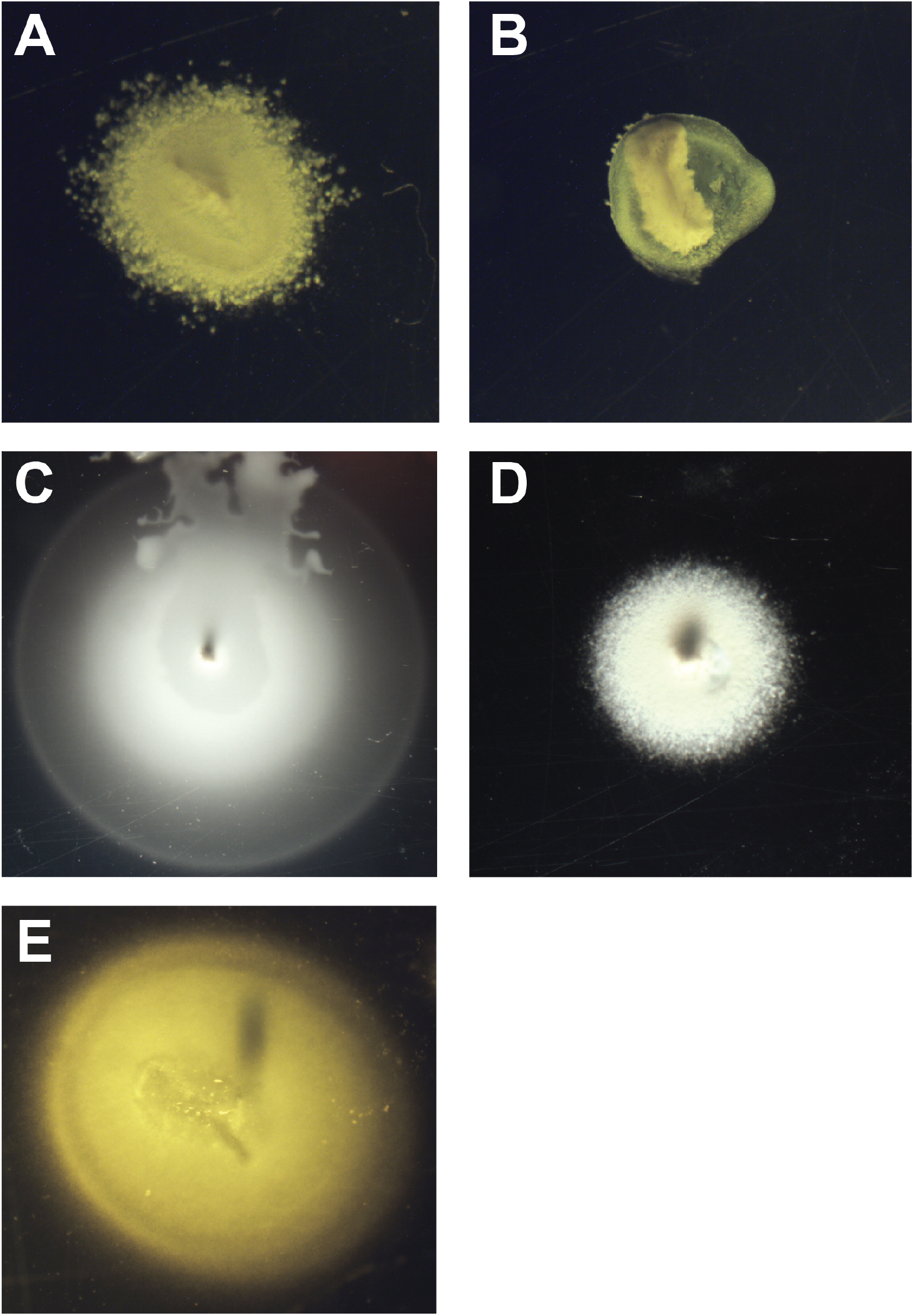
*Psy* B728a exhibiting CS at 30°C (A). A *flgK* mutant (non-motile) incubated in swimming medium at 30°C (B). Spatially separated colonies from a CS plate were picked and transferred to fresh swimming media and incubated at either 20°C (C) or 30°C (D). Normal swimming phenotype exhibited by *Psy* B728a incubated in low-peptone swimming media at 30 °C (E).

We considered the possibility that the CS phenotype might result from a sub-population of cells with gene expression patterns enabling swimming motility that were present in the inoculum, and that prolonged incubation at 30 °C might subsequently suppress motility of all of the cells; such a situation could have allowed a few cells to transiently move beyond the inoculation site until which time as motility was subsequently suppressed at the high temperature, leading them to form colonies at the sites to which they had dispersed. We found, however, that the swimming zone at 30°C continued to expand with additional incubation time and that the constellation phenotype was maintained; the zone always consisted of many punctate, spatially isolated colonies at the periphery of the swimming colony (data not shown). This observation seemed inconsistent with a transiently pre-induced sub-population present only at the beginning of the experiment. These observations suggested transiently motile cells continually arose within the *P. syringae* population incubated at high temperatures, but the majority of the population remained non-motile.

### A subpopulation of cells incubated at 30°C express *fliC* at levels similar to cells incubated at 20°C

The nature of the CS phenotype and the observation that it required a functional flagellum indicated there might be heterogeneity in swimming potential within the population of cells. We hypothesized that such behavioral heterogeneity would also be manifest at the level of flagellin production, and hence expression of *fliC*. Quantitative single cell microscopy confirmed that after 24 hours of incubation at 30°C, a subpopulation cells expressed *fliC* (estimated using a transcriptional reporter) at levels similar to those incubated at 20°C while most cells incubated at the warmer temperature exhibited very low *fliC* expression (Fig. 3). We confirmed this result using flow cytometry (Fig. 4). At 30°C there was a subpopulation of cells that were as fluorescent as the population of cells that were incubated at 20 °C. In this experiment, however, the *fliC*-expressing sub-population was only apparent in cells incubated for 24 hours but not for 12 hours, suggesting that the hyper-expression was dependent on a growth phase of cells seen only in older cultures, perhaps in response to low nutrient conditions that would occur only following replication at the point of inoculation. This result also seemed to support the conclusion above, that the constellation swimming phenotype is not the result of a *fliC-*expressing, motile subpopulation always present in a collection of *P. syringae* cells, but instead, the result of phenotypic heterogeneity that arises only following prolonged incubation at warm temperatures.

**Figure 3.**
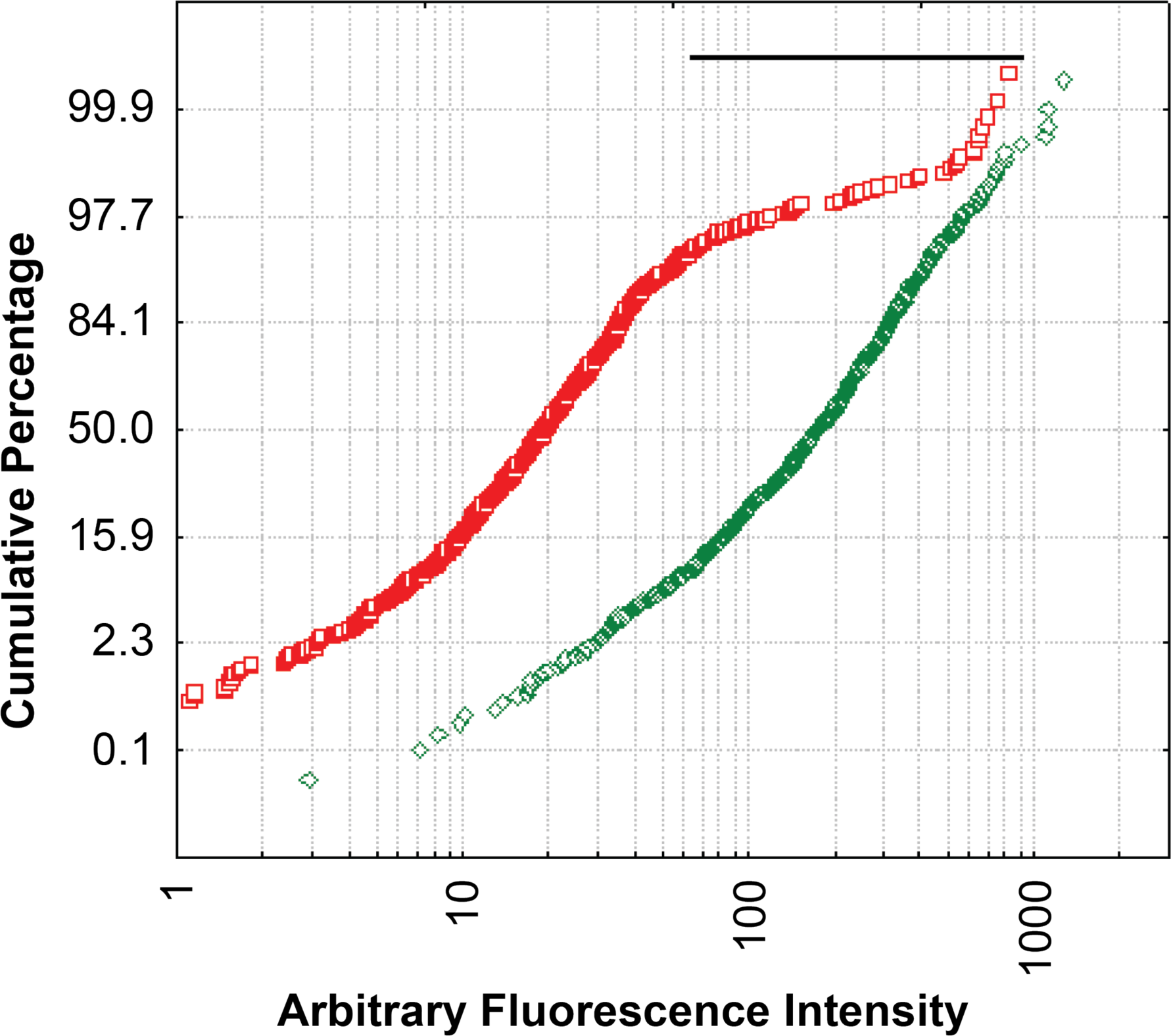
Cumulative normal probability plot of *Psy* B728a harboring a plasmid-borne *fliC* transcriptional reporter (p*fliC*) incubated at either 20°C (green diamonds) or 30°C (red squares) for 24 hours. The subpopulation of cells incubated at 30°C that are is expressing *fliC* at a level higher than expected based on an otherwise normal distribution of GFP fluorescence intensity is indicated under the black line. Experiment was repeated twice with similar results.

**Figure 4.**
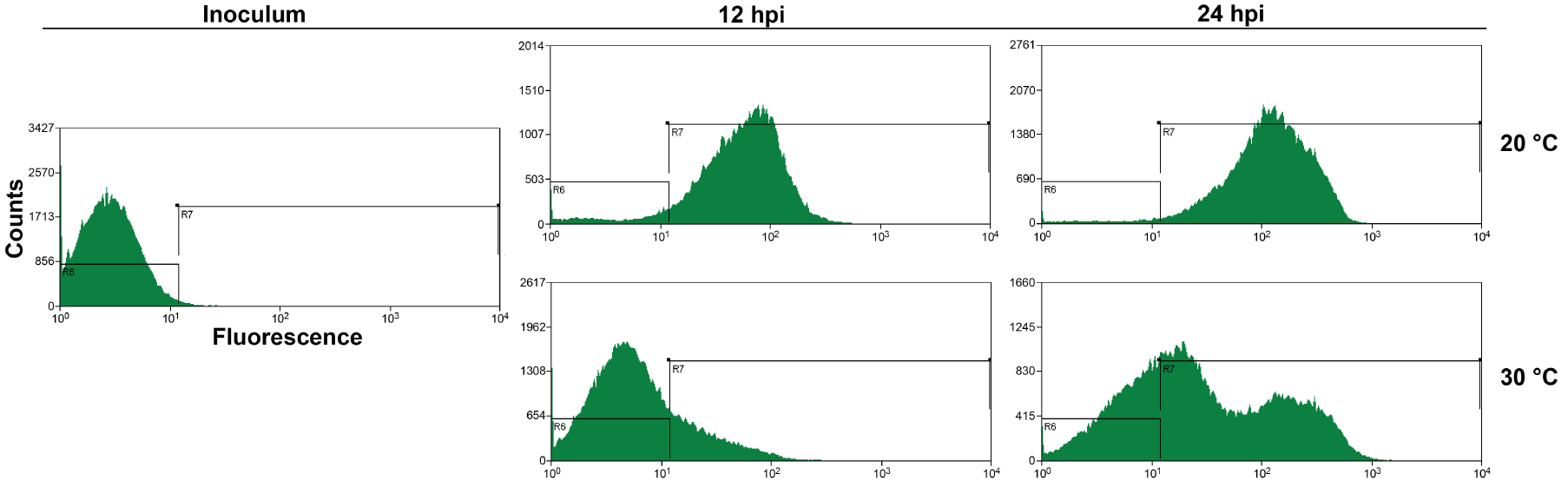
Distribution of GFP fluorescence of *Psy* B728a harboring p*fliC* when incubated at either 20 °C or 30 °C at the indicted times after inoculation. Shown are histograms reflecting the frequency with which cells of a given GFP fluorescence were observed as determined by flow cytometry. Inoculum was incubated over night at 32 °C. Experiment was repeated twice with similar results.

### *P. syringae* exhibits normal swimming at 30°C, as well as elevated *fliC* expression in media with reduced nutrients

As growth phase appeared to contribute to the appearance of a *fliC*-expressing sub-population of cells at 30°C, we tested whether reducing the peptone content of the swimming medium could stimulate motility more uniformly within the population of cells. A 50-fold reduction in peptone concentration resulted in a slow but diffuse swimming phenotype at 30°C (Fig. 2E). Additionally, incubation on media with reduced peptone also resulted in increased *fliC* expression (Fig. 5). Taken together, these results indicate opposing regulation of flagellum-mediated motility, where low nutrients and low temperature stimulate motility, while high temperature and high nutrients suppress motility. Conflicting regulatory signals (e.g. low nutrients and high temperature) result in a heterogeneous population consisting of both motile and non-motile cells, rather than moderately motile cells.

**Figure 5.**
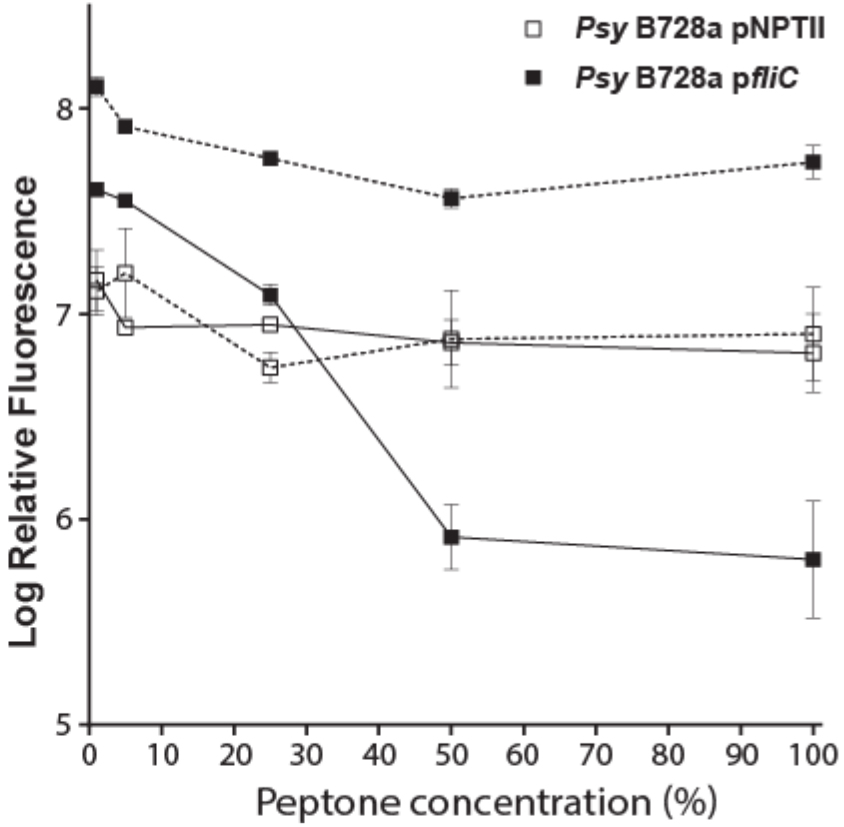
Effect of peptone concentration (as a % of the normal concentration) on *fliC* expression at 20°C and 30°C. See experimental procedures for details for log relative fluorescence calculation. Dashed lines (20°C), solid lines (30°C). Error bars indicate standard deviations with an n of 3 biological replicates. pNPTII is a GFP reporter construct with a temperature insensitive (control) promoter (Hockett et al., 2013).

### Isolation of spontaneous hot swimming (SHS) mutants following prolonged incubation at 30°C

To better understand the genetics underlying transient motility at high temperatures we isolated and characterized stable, hyper-motile mutants that arose spontaneously following several days of incubation at 30°C. Such mutants were identified as flares that developed that phenotypically resembled the normal swimming phenotype exhibited by *P. syringae* at 20°C, as well as increased *fliC* expression (Fig. 6). We thought it likely that these flares consisted of spontaneous mutants that were no longer thermorepressed in motility. Single colonies isolated from the flares were re-inoculated into swimming media at 30 °C. These isolates no longer exhibited the distinctive CS phenotype of the wild type strain, but exhibited the normal diffuse swimming phenotype. Strains recovered in this way were termed spontaneous hot swimming (SHS) mutants. There appeared to be two general phenotypes of the SHS mutants recovered: mutants such as SHS3 that exhibited normal swimming phenotype at both 20°C and 30°C and mutants such as SHS1 that exhibited a swimming defect at 20°C but exhibited normal (non-constellation) swimming at 30°C (Fig. 7 and Table 1). Isolation of SHS mutants in a wild type strain harboring a fusion of a *gfp* reporter gene with the promoter of *fliC* (p*fliC*) on swimming plates at 30 °C revealed that the flares exhibiting a normal swimming phenotype also had relatively high expression of *fliC* compared to the wild type strain in the body of the colony (Fig. 6 top panel). A total of 17 putative mutants that exhibited a shift from a constellation to an enhanced, diffuse swimming phenotype at 30°C were obtained in 3 separate experiments (Table 1).

**Figure 6.**
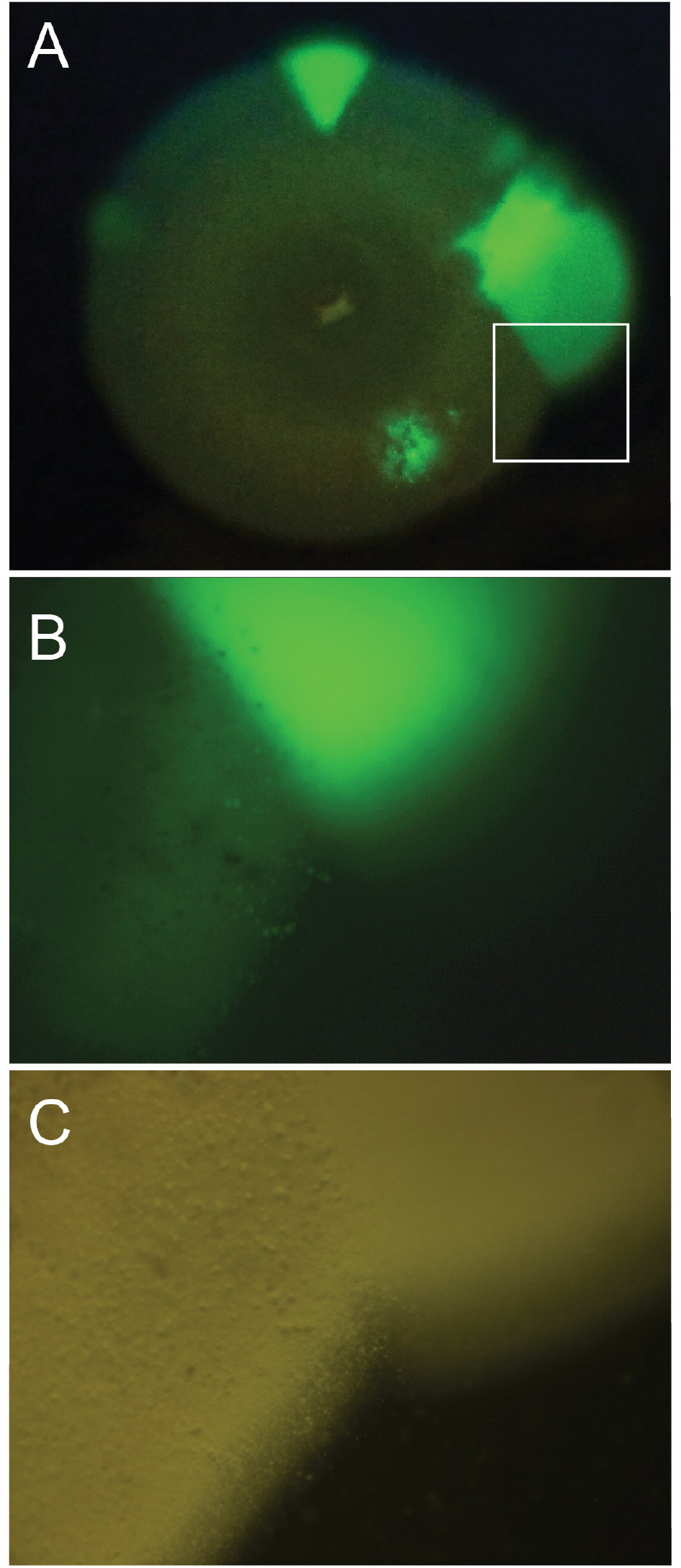
Swimming motility of Psy B728a harboring p*fliC* when incubated at 30°C for 4 days. Green sectors reflect normal swimming (i.e. the swimming phenotype of cells incubated at cooler temperatures) extending from an otherwise CS colony photographed under UV illumination (A). Close up of marked region from top panel (B). Middle panel photographed without UV illumination (C).

**Figure 7.**
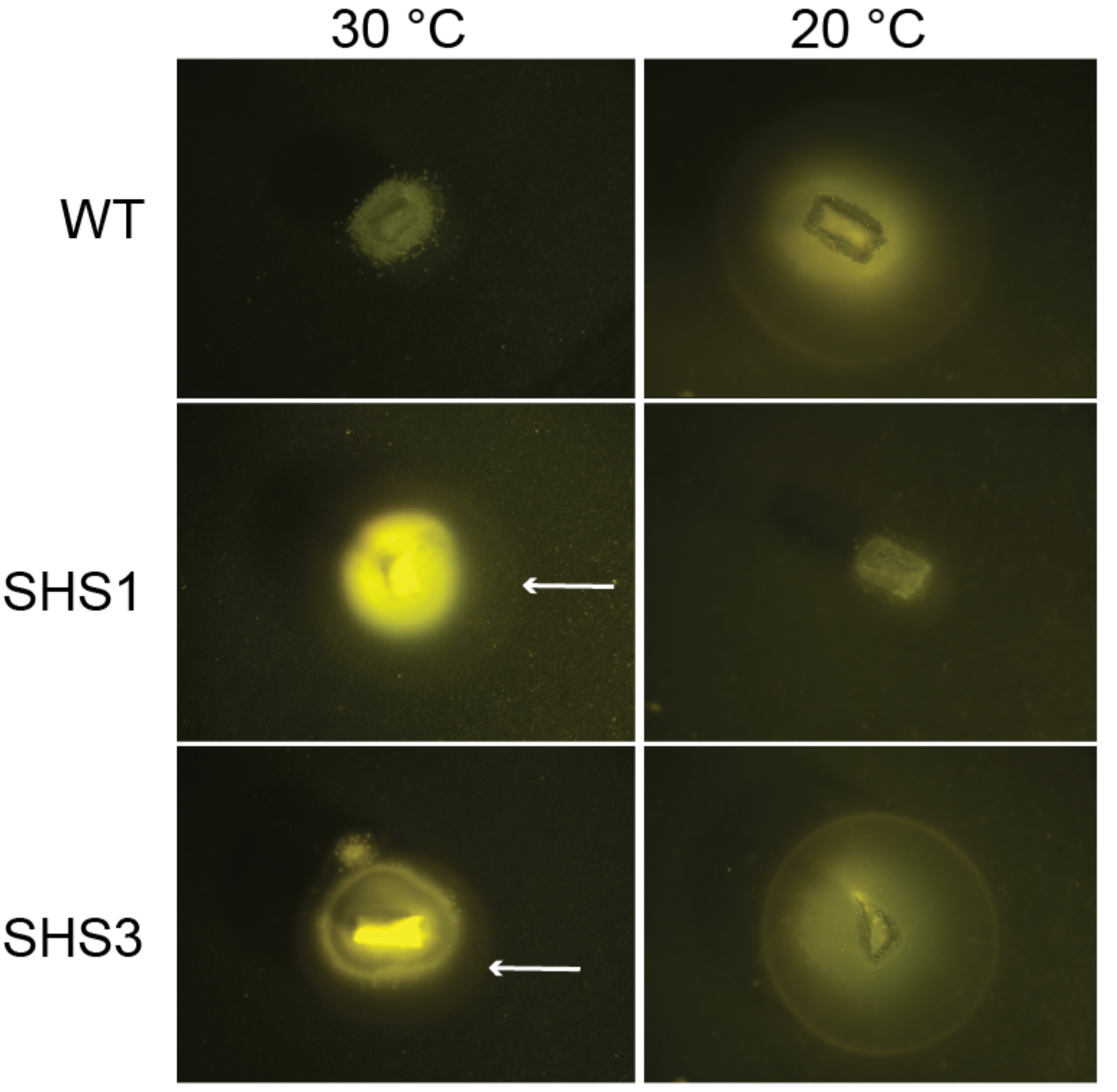
Swimming motility exhibited by *Psy* B728a, as well as SHS mutants 1, and 3, incubated in swimming medium at 30°C (left) or 20°C (right) for 24 hours. White arrows indicate the swimming front within the swimming medium.

**Table 1.** SHS phenotypes observed in different mutants of *Psy* B728a

| strain | isolation | 20 °C swimming phenotype | 30 °C swimming phenotype |
| --- | --- | --- | --- |
| WT | n/a | normal | Constellation |
| SHS1 | 1<sup>st</sup> | reduced | Normal |
| SHS2 | 1<sup>st</sup> | reduced | Normal |
| SHS3 | 1<sup>st</sup> | normal | Normal |
| SHS4 | 1<sup>st</sup> | normal | Normal |
| SHS5 | 1<sup>st</sup> | reduced | Normal |
| SHS6 | 2<sup>nd</sup> | normal | Normal |
| SHS9 | 2<sup>nd</sup> | normal | Normal |
| SHS10 | 2<sup>nd</sup> | normal | Normal |
| SHS13 | 3<sup>rd</sup> | normal | Normal |
| SHS14 | 3<sup>rd</sup> | reduced | Normal |
| SHS15 | 3<sup>rd</sup> | normal | Normal |
| SHS16 | 3<sup>rd</sup> | normal | Normal |
| SHS17 | 3<sup>rd</sup> | reduced | Normal |

In addition to characterizing the swimming phenotypes of the mutants under the two incubation temperatures, we assayed the expression of *fliC* of two different mutants (with distinct mutations, see below) under a range of peptone concentrations at both 20°C and 30°C (Fig. 8). The two mutants (SHS3 and SHS6) both exhibited significantly increased *fliC* expression at 30°C and high peptone concentration compared to the wild type strain. Both mutants also increased *fliC* expression with decreasing peptone concentration, but to a smaller degree relative to the wild type strain.

**Figure 8.**
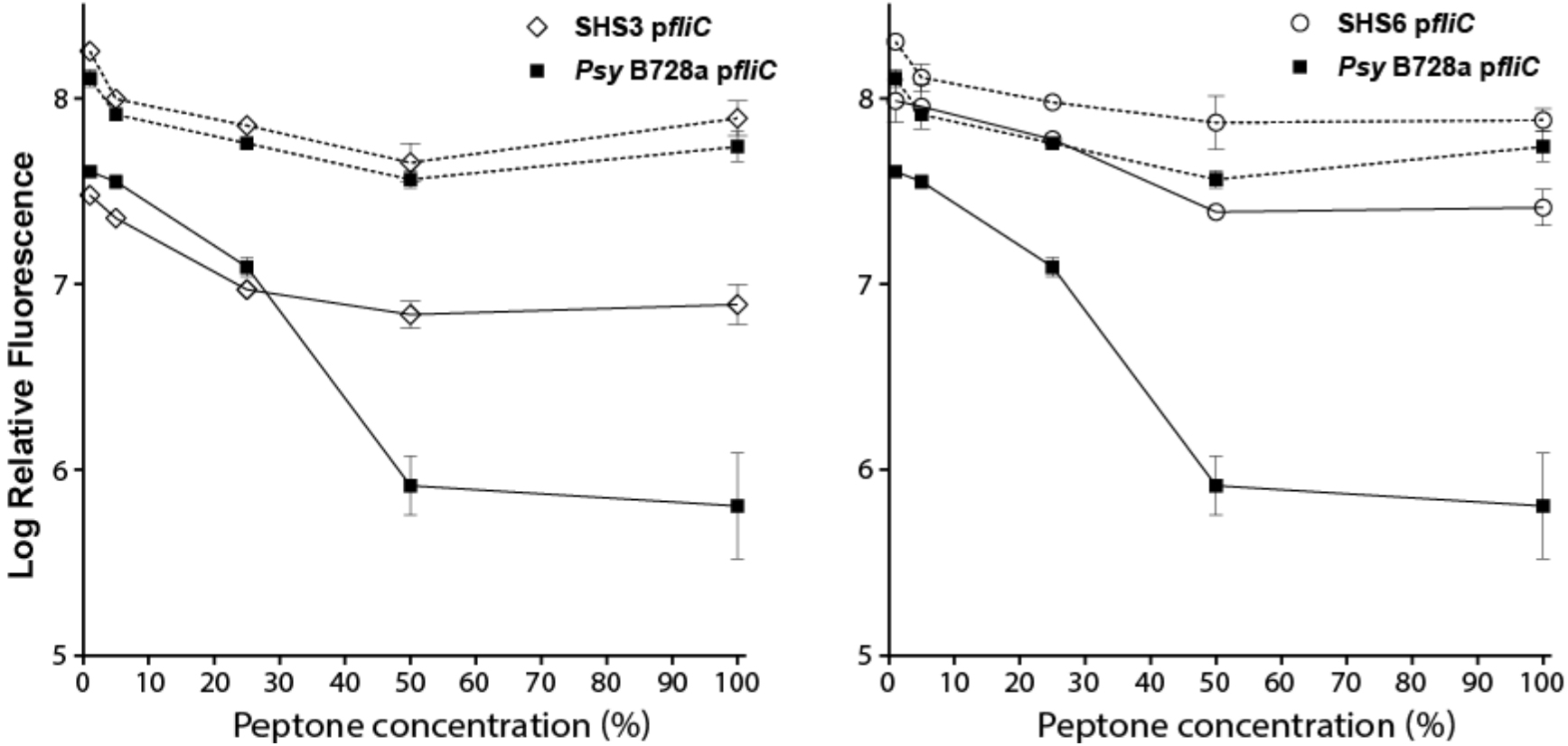
Effect of peptone concentration (as a % of the normal concentration) on *fliC* expression at 20°C and 30°C in *Psy* B728a, SHS3 and SHS6. See experimental procedures for details for log relative fluorescence calculation. Dashed lines (20°C), solid lines (30°C). Error bars indicate standard deviations with an n of 3 biological replicates. Data for *Psy* B728a is identical to that presented in fig. 5, as the data from both figures comes from the same experiment.

### SHS mutants harbor non-synonymous mutations in *fleN*, or in the promoter region of *fleQ*

The genetic lesions in several SHS mutants were identified by comparing the resequenced genomes of these mutants to the wild type ancestor. The most commonly recovered mutations were non-synonomous, single nucleotide polymorphisms (SNPs) in the flagellar gene *fleN* (psyr_3438). Five unique SNPs found in a total of 11 SHS mutants all resulted in a predicted amino acid change in FleN (Table 2). FleN (pseudonym of FlhG in non-Pseudomonads) interacts with the flagellar regulators FleQ and with FlhF, which is encoded by the gene immediately upstream of *fleN* to regulate the number of flagella produced by cells (see discussion). As we had selected mutants that exhibited uniform (non-CS) motility at high growth temperatures, likely a consequence of altered thermoregulation of flagellar synthesis, the abundance of mutations in this gene appeared to confirm our approach to studying heterogeneity in motility at high temperatures. Further analysis of the *fleN* mutants revealed that certain identical SNPs arose repeatedly, such as the A to C transversion harbored by SHS1, 2, 5, 14, and 17, that resulted in a substitution of valine for alanine at position 175.

**Table 2.** Single Nucleotide Polymorphisms observed in SHS mutants of *Psy* B728a

| mutant* | nucleotide position | mutation** | genetic context | amino acid change |
| --- | --- | --- | --- | --- |
| SHS1<sup>a</sup> | 4101623 | A to C | intragenic,<br>psyr_3438 (*fleN*) | V175A |
| SHS2<sup>a</sup> | 4101623 | A to C | intragenic,<br>psyr_3438 (*fleN*) | V175A |
| SHS3<sup>b</sup> | 4101497 | A to T | intragenic,<br>psyr_3438 (*fleN*) | V217E |
| SHS4<sup>b</sup> | 4101497 | A to T | intragenic,<br>psyr_3438 (*fleN*) | V217E |
| SHS5<sup>a</sup> | 4101623 | A to C | intragenic,<br>psyr_3438 (*fleN*) | V175A |
| SHS6 | 4126768 | G to C | intergenic,<br>psyr_3461 psyr_3462 | n/a |
| SHS9 | 4101779 | A to C | intragenic,<br>psyr_3438 (*fleN*) | I123S |
| SHS10 | 4126769 | T to C | intergenic,<br>psyr_3461 psyr_3462 | n/a |
| SHS11 | 4101731 | G to A | intragenic,<br>psyr_3438 (*fleN*) | A139V |
| SHS14 | 4101623 | A to C | intragenic,<br>psyr_3438 (*fleN*) | V175A |
| SHS15 | 4101896 | T to G | intragenic,<br>psyr_3438 (*fleN*) | V84G |
| SHS16 | 4101896 | T to G | intragenic,<br>psyr_3438 (*fleN*) | V84G |
| SHS17 | 4101623 | A to C | intragenic,<br>psyr_3438 (*fleN*) | V175A |

\*mutants sharing the same letter are likely siblings as they were isolated from the same flare
\*\*mutation description is based on positive strand nucleotide

While SHS1, 2, and 5 are likely siblings of the same mutation event because they were recovered from the same flare, SHS 14 and 17 were recovered from different flares on separate plates in a separate experiment. No other SHS mutants isolated with these experiments harbored this particular mutation, indicating that it was not due to an ancestral mutation event present in the founder cells. Likewise, since SHS15 and SHS16, which were isolated from different flares from the same plate, harbored identical mutations they too appear to be a result of independent events.

All five of the unique mutations recovered in *fleN* resulted in a substitution of one aliphatic amino acid for another (A139V, V175A), or a substitution between a non-aliphatic and an aliphatic amino acid (I123S, V84G, V217E). A consequence of these changes is a shift in the size and hydrophobicity of the particular residues, potentially determining whether the residue is buried within the protein or exposed on the surface. Four of the five mutations resulted in reduced hydrophobicity of the amino acid at a site, such as when alanine, glutamic acid, serine, or glycine replaced isoleucine or valine. SHS11 harbored the only mutation that resulted in an increase in residue hydrophobicity. We compared the conservation of these five amino acids across *fleN* homologs in strains where it or its interaction partner *flhF* have been studied (Fig. 9). While none of the particular amino acids were absolutely conserved across all strains examined, in general each residue position usually contained functionally similar residues, usually isoleucine, valine, or alanine, but occasionally methionine. It was noteworthy that the V175A mutation was seen in all five of SHS mutants having the defective cool swimming phenotype (Table 1 and Table 2) while none of the mutants not harboring this mutation exhibited the cool swimming defect.

**Figure 9.**
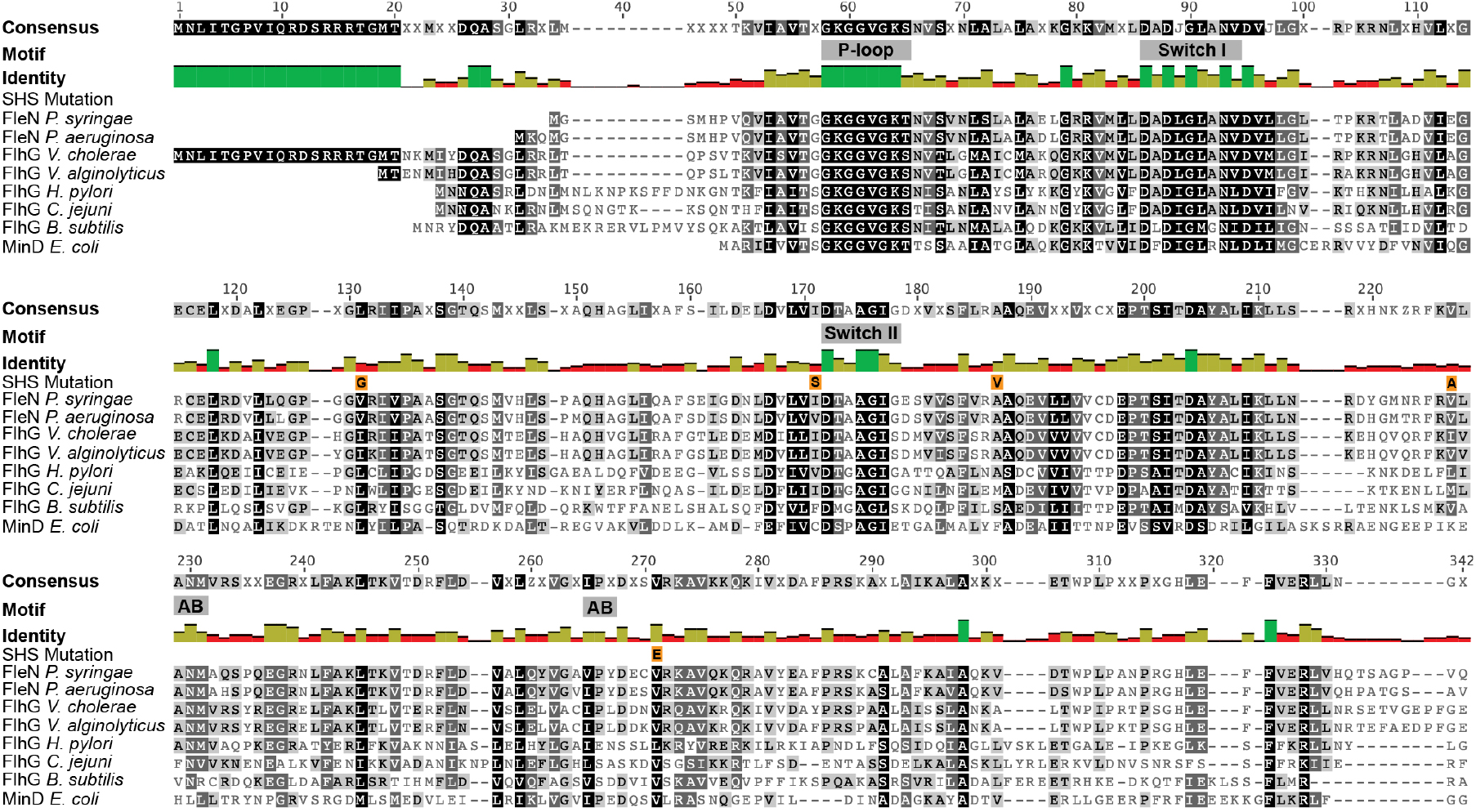
Alignment of *Pseudomonas syringae* FleN with homologs (including the *E. coli* MinD protein) from additional bacterial species. Protein sequence having altered residues in SHS mutants indicated. Motifs described in (Schuhmacher et al., 2015) are shown. AB, ATP binding. Identity graph colors indicate 100% (green), 30-99% (yellow), <30% (red). Shading of residues within the alignment indicates 100% similarity (black), 80-99% similarity (dark gray), 60- 79% similarity (light gray), <60% similarity (unshaded) based on the Blosum62 scoring matrix implemented in Geneious.

SHS6 and SHS10 harbored neighboring SNPs in the promoter region of *fleQ*, the master flagellar regulator (Table 2). Comparison of the *P. syringae fleQ* promoter region with motifs described in its homolog in *P. aeruginosa* (Dasgupta, Ferrell, Kanack, West, & Ramphal, 2002), revealed that neither mutation occurred within the Vfr binding site (Fig. 10). Using BPROM (Solovyev & Salamov, 2011), we found that both mutations occurred within the putative −10 RpoD/σ^70^ binding site. The SHS10 mutation altered the predicted −10 site, while the SHS6 mutation did not (*P. syringae* and SHS10 −10 sites in Fig. 10).

There were 11 SNPs that differed from the sequence of the parental strain that were conserved in all SHS strains (Table S1). Because the identity of these SNPs were conserved across all strains, it seems likely that they accumulated within this particular B728a lineage prior to the mutations that gave rise to the SHS phenotype.

**Figure 10.**
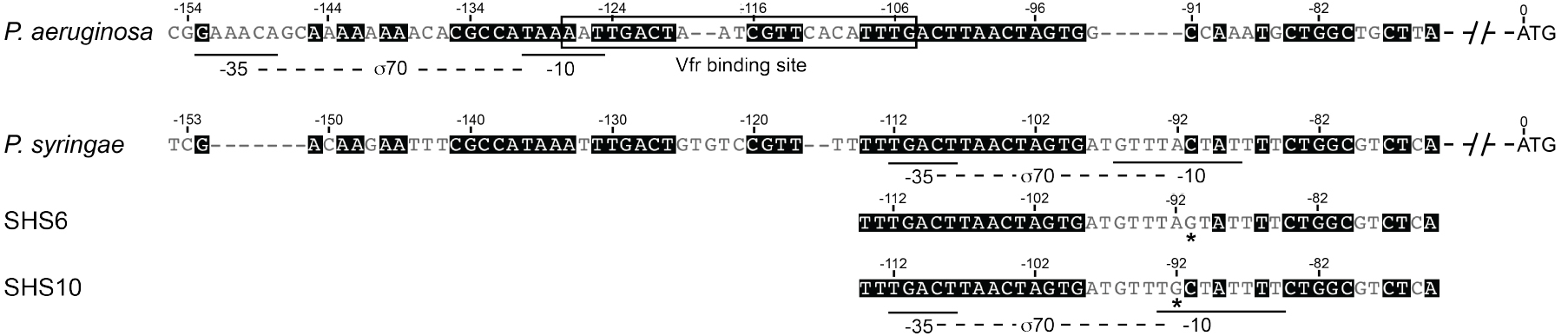
Alignment of the *fleQ* promoter regions of *Pseudomonas syringae* B728a and *Pseudomonas aeruginosa* PA01. *P. aeruginosa* Vfr and σ^70^ binding sites were described in (Dasgupta et al., 2002) while *P. syringae* SHS10 σ^70^ sites were predicted by BPROM (http://linux1.softberry.com/berry.phtml). Asterisks indicate Single Nucleotide Polymorphisms in Spontaneous Hot Swimming mutants.

## DISCUSSION

It is highly significant that FleN was identified here as playing a role in modulating temperature-dependent motility in *P. syringae* as it is central to regulation of motility in other bacterial species. *fleN* (*P. aeruginosa* homolog of *flhG* in *Bacillus*, *Vibrio*, *Helicobacter*, and *Campylobacter* spp.) contributes to several aspects of flagellar regulation. In *P. aeruginosa*, FleN interacts directly with FleQ, the master flagellar regulator in all *Pseudomonas* species thereby inhibiting its ability to stimulate transcription of class II, III, and IV flagellar genes, but it does not affect the expression of *fleQ* or *fliA* themselves (Baraquet & Harwood, 2013; Dasgupta & Ramphal, 2001; Dasgupta et al., 2003; Hickman & Harwood, 2008). In *Vibrio cholera*, FlhG negatively regulates the expression of class II, III, and IV flagellar genes, similar to that in *P. aeruginosa*, but it also negatively regulates the expression of *flrA* (*fleQ* homolog in *Vibrio* spp.) (Correa, Peng, & Klose, 2005). Assuming that FleN in *P. syringae* has a similar effect on flagellar gene regulation as its homolog in *P. aeruginosa* and that modulation of FleN abundance or interaction potential is the determining factor for thermoregulated flagellar gene expression, the previous observations that neither *fleQ* nor *fliA* expression was influenced by incubation temperature would be corroborated (Hockett et al., 2013).

Unfortunately, there is yet no comprehensive understanding of the residues or domains contributing to the interaction of FleN and FleQ. This interaction appears to involve elements in both the N- and C-terminus of FleN, as all truncations tested (including separate C-terminal and N-terminal truncations) resulted in a loss of this interaction (Dasgupta & Ramphal, 2001). ATP binding also appears to be necessary for FleN to interact with FleQ, as a change in the residue predicted to disrupt the ATP binding domain abolished their interaction (Dasgupta & Ramphal, 2001). Although, none of the mutations we recovered in FleN occurred directly within any of the predicted motifs (Fig. 8), some of the SHS mutations may affect the ability of FleN to bind ATP. Additional directed research will be required to address the effect of these mutations on the enzymatic activity of FleN.

In addition to its interaction with FleQ, FleN also physically interacts with another flagellar regulator, FlhF (Kusumoto et al., 2008). FlhF is a signal-recognition particle-type GTPase (as is FleN) that, along with FleN, is found in polarly flagellated bacteria where it is involved in localization of the flagellum, but not in peritrichously flagellated bacteria such as *E. coli* or *Salmonella* spp. (Bange, Petzold, Wild, Parlitz, & Sinning, 2007; Carpenter, Hanlon, & Ordal, 1992; Kusumoto et al., 2008; Kusumoto, Nishioka, Kojima, & Homma, 2009; McCarter, 2004; Pandza et al., 2000). FlhF and FleN reciprocally regulate the number of flagella produced by cells in numerous species. Deletion of FlhF or over-expression of FleN leads to loss of the flagellar filament, while over-expression of FlhF or deletion of FleN leads to the production of multiple polar flagellar filaments (Balaban & Hendrixson, 2011; Correa et al., 2005; Dasgupta, Arora, & Ramphal, 2000; Kusumoto et al., 2008; 2009; 2006). Multi-flagellation is associated with reduced swimming ability, an effect that was clearly observed in SHS1, 2, 5, 14, and 17 at 20°C. Presumably, the multiple flagellar filaments form bundles that cannot function as well as single filaments. In general, those SHS mutants that did not exhibit a swimming defect at cool temperatures exhibited stronger swimming at 20°C than at 30°C. This indicates that while these mutants are capable of normal swimming at 30°C (as opposed to constellation swimming), they are somewhat defective in flagellum formation, or some other aspect of swimming at 30°C.

Whether temperature directly influences the structure of FleN is currently unknown. Based on our finding of SNPs primarily in *fleN* in the SHS mutants, we hypothesize that either the overall structure of FleN, or that of a particular FleN domain, is temperature sensitive. While we hypothesize that FleN is the component governing thermal suppression of motility in *P. syringae*, implying that FleN directly senses and responds to temperature, the alternative hypothesis can be formulated that FleN expression itself is thermo-regulated, being more abundant at 30°C than at 20°C. If FleN has increased expression at 30°C, this is apparently not due to its elevated transcription since similar amounts of *fleN* messages were found in cells grown at different temperatures (Hockett et al., 2013). It is possible that *fleN* is translated more efficiently at 30°C. If FleN is indeed more abundant at 30°C than 20°C, then the SHS mutants with altered FleN that are not defective in swimming at 20°C may exhibit a temperature-sensitive phenotype, resulting in selective loss of FleN activity at 30°C, but not at 20°C. Such temperature-sensitive mutations commonly result from single amino acid substitutions that alter the folding, and thus the function, of the protein at the non-permissive temperature (Baer, Wesolowski, & Altman, 1989; Goldenberg & King, 1981; Nguyen Truong Hoai Thu, Pratt, Rule, Hsue, & Ho, 1991). It is possible that the V175A substitution in SHS mutants having a swimming defect at 20°C, conferred a more pronounced change in FleN folding such that it was non-functional even at 20°C. Finally, it remains possible that FleN is neither itself sensitive to temperature, nor is its expression temperature-dependent, but that the mutations in *fleN* negate the effects of an alternative and currently unknown thermoregulator. In any event, the *fleN* mutations recovered in this work should prove valuable in further investigation of the physical interaction of FleN with flagellar regulatory components such as FleQ and FlhF, as well as the potential effect of temperature on these interactions.

While the majority of SHS mutants supported the role of FleN in thermoregulation of motility, circumstantial evidence also suggests that other flagellar components also play a role in this process. The two SHS mutants with SNPs in the *fleQ* promoter region are intriguing based on their proximity to each other as well as their co-localization with a predicted σ^70^ binding site. In *P. aeruginosa* a Vfr binding site overlaps the σ^70^ −10 site (Dasgupta et al., 2002) (Fig. 9). Vfr is a homolog of CAP (also known as CRP, cAMP receptor protein) in *E. coli;* cAMP being a central mediator of catabolite repression (Botsford & Harman, 1992). In *E. coli*, flagellar synthesis, and thus motility, requires CAP to be bound to cAMP, and thus is catabolite repressible (Adler & Templeton, 1967). The expression of the *flhDC* operon is also subject to catabolite repression (Soutourina et al., 1999). Certain spontaneous mutants in *E. coli* that no longer exhibit repressed flagella synthesis at high temperatures also have other pleiotropic defects such as a lack of catabolite repression, while some other spontaneous mutants lacking such catabolite repression are also able to synthesize some flagella at elevated temperatures, indicating either that there is a functional link or common regulator between catabolite repression and thermoregulation in flagella synthesis in *E. coli* (Li, Louise, Shi, & Adler, 1993; Silverman & Simon, 1974; 1977). The contribution of Vfr to flagellar regulation in *Pseudomonas* is poorly understood, and may differ from that in *E. coli*. Over-expression of Vfr in *P. aeruginosa* results in reduced expression of *fleQ* as well as reduced motility (Dasgupta et al., 2002; Silverman & Simon, 1974). Currently, we have two lines of evidence suggesting that nutrient status regulates *fliC* expression and motility in *P. syringae* (Figs. 2 and 5). First, the sub-population of cells in wild type cells that highly expressed *fliC* even at 30°C only accumulated after a visible colony had formed. Secondly, reduction of the peptone concentration in the medium (the major nutrient source, particularly for N) enabled a normal (diffuse) swimming colony phenotype and increased *fliC* expression even when incubated at 30°C. We speculate that upon the formation of a visible colony there would have been sufficient cell growth to deplete some key nutrient(s) in the local environment that otherwise would have repressed *fliC* expression at 30 °C. Greatly reducing the peptone concentration in the media itself presumably mimics the release from nutrient repression of movement that otherwise would have occurred only near colonies. More study will be needed to better describe the putative linkage between catabolite repression and thermoregulation of flagella synthesis and motility in *P. syringae*. It is noteworthy that *vfr* (psyr_4577) was stimulated under cool incubation conditions in *P. syringae* (Hockett et al., 2013) indicating that if it does bind to the *fleQ* promoter in *P. syringae* (which is currently unknown) then it probably does not inhibit its expression. We previously showed that neither *fleQ* nor *fliA* (the two putative class I regulators of flagellar synthesis in *P. syringae* are thermoregulated at the level of transcription (Hockett et al., 2013). Based on the presence of the SNPs of two SHS mutants in the promoter region of *fleQ*, we would expect that these mutations would affect its transcription. These results suggest that, although repression of *fleQ* transcription at high temperatures cannot be the determinant of suppression of flagellar production at high temperatures, over-expression of *fleQ* may be sufficient to mask down-stream thermoregulation, potentially by over-expressing *flhF* compared to *fleN*.

Heterogeneity in flagellin expression has been observed in animal pathogens, such as *E. coli* and *Salmonella enterica* serovar Typhimurium (Freed et al., 2008; M. K. Stewart, Cummings, Johnson, Berezow, & Cookson, 2011). In *Salmonella*, heterogeneous expression of flagellin aids in evasion of the caspase-1 inflammatory response (M. K. Stewart et al., 2011). Heterogeneous flagellin expression *in vitro* is controlled by YdiV, a protein with weak homology to c-di-GMP-degrading EAL domain-containing proteins. Intriguingly, YdiV acts by binding to FlhD_4_C_2_ mediating both stochastic as well as deterministic flagellin expression in response to nutritional cues (Koirala et al., 2014; Wada et al., 2011). Thus similarities to the processes seen to enable heterogeneous motility in *P. syringae* at high temperatures are apparent.

The phyllosphere is characterized by rapid shifts in environmental conditions, such as temperature and water availability (Hirano & Upper, 2000a; Lindow & Brandl, 2003). While the duration of water availability is often inversely related to temperature (water is more persistent or likely under conditions of cool temperatures associated with night or rainfall events), it is not a guarantee that an elevated temperature will always lead to reduced water availability. In the event that water availability coincides with elevated temperatures, it may be beneficial for the bacterial population to contain a sub-population of cells that are capable of exploring/exploiting its local habitat. Such exploration would be facilitated by active motility. For example, if water did become sufficiently available even under warm conditions, at least these few cells could explore the leaf surface and exploit un-colonized sites having nutrient abundance. A pre-existing sub-population that was capable of motility because of a lack of catabolite repression, and despite the occurrence of high temperatures could then colonize the leaf during periods where water is not sufficiently available long enough to elaborate the flagellum and chemotaxis system de novo. Such cells could thus be thought of as exceptions to an otherwise conservative strategy of colonization that avoids movement except under cool conditions, when motility will have a higher probability of being successful. The mutants uncovered in this study should prove valuable as tools for investigating how different patterns of thermoregulated motility as well as heterogeneous motility of *P. syringae* affect its fitness on plants.

## EXPERIMENTAL PROCEDURES

### Bacterial strains, plasmids, culture media, and growth conditions

*Pseudomonas syringae* pv. *syringae* B728a (*Psy* B728a) (Loper & LINDOW, 1987) and derivative strains were routinely cultured in King’s medium B (KB) broth, or on KB plates supplemented with 1.5% (w/v) Difco agar technical (BD, Sparks, MD) at 28°C (E. O. King, Ward, & Raney, 1954), except for experiments where different incubation temperatures are noted. Antibiotics were used at the following concentrations: gentamicin, 15 μg/mL; rifampicin, 100 μg/mL.

Temperature-dependent assays were performed in incubators adjusted to the appropriate temperature as described in the text. Plate temperatures were routinely monitored using a CZ-IR thermometer (ThermoWorks, Lindon, UT). Incubator temperatures and RH were routinely monitored using HOBO data loggers (Onset, Bourne, MA).

### Swimming assays

Swimming media was prepared as described in (Hockett et al., 2013), except in the low peptone swimming media, where only 1.0 % of the normal concentration of peptone was added (0.2 g peptone/L; all other media components were added at 50% of normal concentration).

#### Transcriptional reporter assays

Strains were grown overnight in KB media with appropriate antibiotic. Cultures were washed twice with phosphate buffer (pH 7.0) and resuspended to ~OD_600_ = 0.1. Five microliters of each strain was spotted in triplicate onto KB plates with varying concentrations of proteose peptone #3 (BD Biosciences) to simulate starvation conditions and incubated for 24 hours. Each colony was recovered by taking an agar core using a cork borer and sonicated for 10 min in phosphate buffer. The sonicated colonies were vortexed to mix, and were transferred to black clear flat bottom polystyrene untreated microplates (Corning #3631). Each colony was assayed in triplicate, resulting in nine independent measurements per strain per treatment. Fluorescent limits were set to 480 nm excitation and 510 nm emission with bandwidths of 9 nm and 15 nm for emission and excitation, respectively, with a Spectramax i3x platereader (Molecular Devices.) Absorbance was read at 600 nm with a water constant pathcheck. Background fluorescence and optical density was subtracted by preparing cores without colonies and processing them exactly as samples. Relative fluorescence was determined by dividing the arbitrary fluorescence values by optical density for each well.

### Quantitative microscopy and flow cytometry

Quantitative single cell microscopy was performed on cells that were grown for 24 hours at various temperatures on KB amended with gentamicin. Following incubation, cells were fixed in 2 % formaldehyde (w/v, final concentration) and stained with DAPI (2.5 μg/μL, final concentration) for 20 minutes. Cells were observed with a Zeiss AxioImager fluorescence microscope with DAPI (325 to 375 nm band pass excitation, and 460 nm long pass emission) and GFP (450 to 490 nm band pass excitation, and 500 to 550 nm band pass emission filter) filter sets (Chroma; Bellows Falls, VT). Images were captured using a Hamamatsu Orca 03 charge coupled device (CCD) camera. The intensity of GFP fluorescence of individual cells was determined using iVision-Mac software. Cumulative probability plots of cell GFP intensities were generated using Statistica (StatSoft; Tulsa, OK).

Cells were similarly prepared (except not stained with DAPI) for flow cytometry, which was performed with a Cytopeia INFLUX Sorter using a 488 nm excitation laser and a GFP detector. Gating was based on cell size, determined by forward scatter and side scatter where significant counts occurred in cell-containing samples but not in buffer controls. Data was plotted with Summit v3.1 (Cytomation, Inc; Fort Collins, CO).

### Spontaneous Hot Swimming mutant isolation

Swimming plates were incubated at 30°C for 3 to 4 days, until robust flares had formed that resembled the normal, diffuse swimming phenotype. A culture from the tip of each flare was streaked onto KB plates to enable isolation of single colonies. These single colonies were used to inoculate fresh swimming medium which was incubated at 30°C or 20°C. *Psy* B728a was routinely included for phenotype comparison. Colonies that exhibited the normal swimming phenotype at 30°C (i.e. non-constellation), regardless of their 20°C phenotype, were stored at −80 °C.

### Sequencing of Spontaneous Hot Swimming mutants

Genomic DNA was isolated from SHS mutants grown overnight in KB broth using a DNeasy Blood and Tissue Kit (QIAGEN; Valencia, CA). Samples were treated with 10 μl RNAse A (10 mg/ml, Fermentas; Glen Burnie, MD) to remove RNA contamination. Isolated genomic DNA was fragmented with a Covaris-S220 adjusted for 300 base pair fragments according to the manufacturer’s protocol (Covaris, Inc; Wourn, MA). Genomic fragments were end-repaired and A-tailed similar to cDNA samples described in (Hockett et al., 2013). DNA Samples were routinely purified using AMPure XP beads (Beckman Coulter, Brae, CA). Illumina adapters were ligated onto fragments as described in (Hockett et al., 2013) and fragment sizes between 300 and 600 nucleotides were selected using various AMPure XP bead-to-sample ratios, which were empirically determined to recover the desired fragment sizes as determined using a 100 bp ladder (Life Technologies; Carlsbad, CA). Following fragment sizing, samples were amplified with bar-coded Illumina primers. Amplified libraries were quantified using a Qbit dsDNA HS assay (Life Technologies) and fragment size distribution was determined with a 2100 bioanalyzer (Agilent; Santa Clara, CA). Sequencing consisted of 100 bp paired-end reads, from a single multiplexed lane using an Illumina HiSeq 2000 at the Vincent J. Coates Genomics Sequencing Laboratory at the University of California, Berkeley.

Reads were aligned to the *P. syringae* B728a genome using bowtie 0.12.7 (Langmead, Trapnell, Pop, & Salzberg, 2009) to identify single nucleotide polymorphisms (SNP) and allowing for a maximum of three SNPs in a given read. Aligned reads were subsequently processed using the SNP detection feature on CLC Genomics Workbench (CLC Bio; Cambridge, MA) with default settings. Predicted SNPs were considered well supported if greater than 80 % of the mapped reads predicted an identical SNP.

The following FleN homologs were aligned in Geneious version 6.1.8 (Kearse et al., 2012) using the MAFFT alignment option (G-INS-i algorithm, BLOSUM30 scoring matrix, with a 1.53 gap open penalty, and 0.123 offset value): *Escherichia coli* MinD (accession ABC70502.1), *Vibrio cholerae* FlhG (accession NP_231699.1), *Vibrio alginolyticus* FlhG (accession BAD93655.1), *Helicobacter pylori* FlhG (accession CBI66709.1), *Campylobacter jejuni* FlhG (accession ALF92767.1), *Bacillus subtilis* FlhG (accession AIY97254.1), *Pseudomonas aeruginosa* PAO1 FleN (locus tag PA1454, accession NP_250145.1), and *Pseudomonas syringae* B728a FleN (locus tag psyr_3438, accession YP_236508.1).

The *fleQ* promoter regions from *P. aeruginosa* PAO1 and *P. syringae* B728a were aligned in Geneious using the MAFFT alignment option (E-INS-I algorithm, 200PAM/k=2 scoring matrix, with a 2 gap open penalty, and 0 offset value).

### Promoter prediction

*fleQ* promoter site prediction was performed with the Virtual Footprint promoter analysis option, which included all *P. aeruginosa* and *E. coli* position weight matrices (Münch et al., 2005). Binding sites were predicted using BPROM (available at http://linux1.softberry.com/berry.phtml).

## ACKNOWLEDGEMENTS

We thank Brian Staskawicz for use of CLC Genomics Workbench software and computing cluster. We thank Hector Nolla for the assistance with the flow cytometry experimentation and data analysis.

We thank Stacey Wood for assistance in quantifying the combinatorial effect of temperature and nutrient levels on *fliC* expression.

## AUTHOR CONTRIBUTIONS

Study conception and design: KLH and SEL. Performed experiments and analyzed data: KLH. Contributed to writing the paper: KLH and SEL.

---

**Table S1.** Single Nucleotide Polymorphisms common to all sequenced SHS mutants of *Pseudomonas syringae*

| location | mutation* |
| --- | --- |
| 376070 | A to G |
| 376071 | G to A |
| 739416 | T to C |
| 739417 | C to G |
| 739465 | T to G |
| 951716 | C to A |
| 951765 | C to G |
| 951817 | C to G |
| 951882 | C to G |
| 5423688 | C to T |
| 5426993 | T to C |

\*mutation designated based on sense strand

**Table S2.** Bacterial strains and plasmids.

| Strain or plasmid | Relevant characteristics | Source or reference |
| --- | --- | --- |
| *P. syringae* strains |  |  |
| *P. syringae* B728a | Wild type, Rf<sup>R</sup> | (48) |
| $\Delta flgK$ | *P. syringae* B728a harboring a *flgK* deletion | (8) |
| SHS1 | Non-CS swimmer at 30°C derivative of *P. syringae* B728a | This work |
| SHS2 | Non-CS swimmer at 30°C derivative of *P. syringae* B728a | This work |
| SHS3 | Non-CS swimmer at 30°C derivative of *P. syringae* B728a | This work |
| SHS4 | Non-CS swimmer at 30°C derivative of *P. syringae* B728a | This work |
| SHS5 | Non-CS swimmer at 30°C derivative of *P. syringae* B728a | This work |
| SHS6 | Non-CS swimmer at 30°C derivative of *P. syringae* B728a | This work |
| SHS9 | Non-CS swimmer at 30°C derivative of *P. syringae* B728a | This work |
| SHS10 | Non-CS swimmer at 30°C derivative of *P. syringae* B728a | This work |
| SHS11 | Non-CS swimmer at 30°C derivative of *P. syringae* B728a | This work |
| SHS14 | Non-CS swimmer at 30°C derivative of *P. syringae* B728a | This work |
| SHS15 | Non-CS swimmer at 30°C derivative of *P. syringae* B728a | This work |
| SHS16 | Non-CS swimmer at 30°C derivative of *P. syringae* B728a | This work |
| SHS17 | Non-CS swimmer at 30°C derivative of *P. syringae* B728a | This work |
| Plasmids |  |  |
| *pfliC* | pPROBE-GT harboring the *fliC* promoter, Gm<sup>R</sup> | (54) |

Rf, rifampicin; Gm, gentamicin

